# Quantum analysis of protein–ligand binding by integrating structural resolution, sequence homology, and ligand properties

**DOI:** 10.1101/2025.06.27.661905

**Authors:** Don Roosan, Samira Samrose, Rubayat Khan, Saif Nirzhor, Brian Provencher

**Affiliations:** School of Engineering and Computational Sciences, Merrimack College, 315 Turnpike St, North Andover, MA 01845, United States; Utah State University, 4205 Old Main Hl, Logan, UT 84322, United States; University of Nebraska Medical Center, S 42nd &, Emile St, Omaha, NE 68198, United States; University of Texas Southwestern Medical Center, 5323 Harry Hines Blvd, Dallas, TX 75390, United States; Department of Natural Sciences, Merrimack College, 315 Turnpike St, North Andover, MA 01845, United States

## Abstract

Predicting protein–ligand binding affinity is a fundamental challenge in computational biology and drug discovery, complicated by diverse factors including protein sequence variability, ligand chemical diversity, and structural resolution. Here, we present an integrative study that combines classical machine learning and quantum-enhanced modeling to investigate how crystal structure resolution, sequence similarity, and ligand properties jointly influence binding affinity. Using a curated “refined” dataset from PDBbind and an expanded general dataset, we first conduct correlation and regression analyses to quantify the relationships among binding affinity, ligand descriptors (e.g., molecular weight, logP), and protein structural metrics (resolution, R-factor). We observe moderate positive correlations between ligand size/hydrophobicity and affinity, and a slight negative correlation between resolution and affinity in the refined dataset that largely disappears in the general set. We then train multiple predictive models, including random forests, deep neural networks, and quantum-enhanced approaches—quantum kernel methods, variational quantum circuits, and a hybrid classical–quantum neural network. Experimental results show that quantum-enhanced models perform on par with classical methods in predicting binding affinities and, in some cases, offer modest improvements. Notably, a hybrid quantum–classical model achieves the highest accuracy (Pearson correlation R≈0.80R) on the refined dataset. These findings highlight the potential of quantum computing for capturing complex patterns in biomolecular data, laying groundwork for improved structure-based drug design. Our study underscores that while data quality and curation greatly influence observed trends, quantum machine learning despite current hardware limitations can already serve as a competitive and promising tool in computational structural biology.

**AUTHOR SUMMARY:** In our study, we confront a major bottleneck in the creation of new medicines: accurately predicting how strongly a potential drug will attach to its target protein in the body. Getting this right early in the process could save billions of dollars and years of research by preventing dead-ends. We investigated if quantum computing, a new technology that processes information in a fundamentally different way, could provide a better solution. We trained several computer models to predict these binding strengths, comparing standard artificial intelligence (AI) with new models enhanced by quantum machine learning. Our results showed that a hybrid model, which strategically combines classical AI with a quantum-powered component, delivered the most accurate predictions on high-quality, curated data. This work demonstrates that quantum computing is already a competitive tool for real-world biological problems, not just a future possibility. We believe that by further developing these quantum-enhanced approaches, we can create more reliable predictive tools to make the long and difficult search for new drugs faster and more successful.

## INTRODUCTION

Protein–ligand interactions lie at the heart of biochemical processes and drug action, yet they are remarkably complex. When a small molecule binds to a protein, a delicate balance of forces— hydrogen bonding, hydrophobic effects, electrostatic interactions, and conformational changes—determines the binding affinity. Capturing this interplay is challenging because even minor structural or chemical changes can shift the interaction landscape. As a result, understanding and accurately predicting binding affinity for protein–ligand complexes remains a major challenge in computational biology and chemistry.

This challenge has profound implications for drug discovery. Designing effective therapeutics hinges on finding molecules that bind strongly and specifically to disease-related protein targets. However, the drug discovery pipeline is notoriously inefficient: only a tiny fraction of screened compounds ever become approved drugs, and many candidates fail in late-stage trials due to insufficient efficacy or unforeseen interactions [1], [2]. These failures come at a tremendous cost, with the development of a single successful drug often taking over a decade and costing billions of dollars. Improving early predictions of binding affinity could help identify promising drug candidates faster and filter out weak binders, thereby saving time, reducing costly late-stage failures, and streamlining the development process. There is, therefore, a pressing need for better computational models that can reliably predict protein–ligand binding outcomes.

A key reason drug discovery is so difficult is the multitude of factors that influence protein–ligand binding affinity. Molecular size and shape determine whether a ligand can snugly fit into the protein’s binding pocket. Hydrophobicity and polarity play crucial roles: hydrophobic regions of a drug tend to seek out hydrophobic pockets on the protein, while polar or charged groups may form specific hydrogen bonds or salt bridges that stabilize the complex [3]. The protein’s flexibility and the structural resolution of available protein-ligand complex data further affect our ability to model interactions—high-resolution structures allow detailed mapping of contacts, whereas low-resolution or unknown binding conformations add uncertainty [4]. Accounting for all these factors simultaneously is non-trivial. Often, trade-offs arise (for example, a chemical modification might improve a hydrogen bond but reduce hydrophobic complementarity), making the accurate prediction of binding affinity an intricate multi-dimensional problem.

Given the complexity, researchers have increasingly turned to machine learning (ML) to improve binding affinity prediction. Data-driven models can learn complex patterns from large databases of protein–ligand complexes, capturing subtle relationships that may elude human-designed scoring functions. Indeed, modern ML approaches, including deep neural networks and other regression/classification models [5], [6], [7], [8], [9], have achieved better predictive accuracy than many classical physics-based methods for affinity prediction [10]. These models have become valuable tools in computer-aided drug design, where they can rapidly screen virtual compounds and prioritize those likely to bind strongly [11]. However, challenges remain: conventional ML models often struggle to generalize beyond the specific proteins and chemotypes present in their training data. An ML model trained on one set of protein families or molecular scaffolds may perform poorly when confronted with novel targets or chemistries [12]. Moreover, achieving high accuracy can require extremely large models and datasets, which leads to high computational costs for training and hyperparameter tuning. This combination of generalizability issues and computational overhead means that, despite progress, classical ML is not a panacea for binding affinity prediction [13], [14], [15], [16], [17].

Quantum machine learning (QML) has recently emerged as a promising avenue to address some of these challenges. QML integrates quantum computing principles into machine learning algorithms, offering fundamentally new ways to represent and process data. One notable advantage of QML is its ability to perform high-dimensional feature mapping naturally: a quantum computer can encode input data (such as molecular descriptors or even raw molecular structures) into quantum states in a Hilbert space of exponentially large dimension [18]. In essence, quantum models can explore extremely rich feature spaces, enabling the detection of patterns or correlations that might be inaccessible to classical models working in lower-dimensional spaces. Additionally, quantum algorithms exploit phenomena like superposition and entanglement, which allow them to recognize complex nonlinear patterns with a different computational paradigm [12]. This means a well-designed QML model might capture the intricate, multi-body relationships in protein–ligand data (such as subtle cooperative effects between different parts of a molecule and a protein) more effectively or efficiently than a classical model. In theory, QML approaches could thus improve the accuracy of binding affinity predictions or achieve similar accuracy with less data by leveraging quantum-enhanced pattern recognition.

In particular, several quantum-enhanced machine learning methods are being explored for their potential impact on problems like binding affinity prediction: Quantum Support Vector Machines (QSVMs) use quantum computing to implement kernel-based learning in extremely high-dimensional spaces. By encoding molecular features into a quantum state, a QSVM can compute complex similarity measures between data points that would be intractable classically. This approach can enhance the ability to separate and predict outcomes based on subtle feature differences, potentially leading to more accurate classification or regression models for binding affinity.

A Variational Quantum Circuit (VQC) is a type of quantum neural network where a parameterized quantum circuit is trained to learn a given task. Using classical optimization to adjust quantum gate parameters, VQC models can approximate complicated functions and decision boundaries. In the context of binding affinity, a variational quantum circuit can be designed to take in molecular input data and output a predicted affinity, with the quantum circuit’s entangled qubits representing complex feature combinations [19]. Such models leverage quantum superposition to compactly represent relationships, and early studies indicate they can learn patterns in chemical data, all while being run on near-term quantum hardware.

Because current quantum computers are limited in size and noise, practical QML algorithms often use a hybrid approach. In a hybrid model, classical and quantum computations are interwoven, each doing what it does best. For example, a classical computer might handle data preprocessing and overall control flow, while a quantum processor is tasked with solving a sub-problem like evaluating a complex molecular kernel or optimizing part of a model’s parameters through quantum circuits [20]. This synergy allows the utilization of quantum advantages on key bottlenecks (such as exploring a vast chemical feature space) without requiring a fully quantum solution. Hybrid models have the potential to accelerate certain computations or improve model quality, and importantly, they can be tested on today’s quantum machines by offloading the heavy lifting to quantum circuits and using classical routines to support and interpret the results.

Motivated by these developments, the objective of this paper is to develop and assess quantum-enhanced machine learning models for protein–ligand binding affinity prediction. We aim to demonstrate the feasibility of applying QML techniques to real-world binding affinity data and to evaluate their performance relative to state-of-the-art classical ML approaches. By doing so, we seek to highlight the practical advantages that quantum machine learning can bring to drug discovery, such as improved prediction accuracy or efficiency, and to provide insights into how these quantum approaches can be integrated into future computational drug design pipelines. In summary, this work explores how quantum-enhanced models can better capture the complexities of protein–ligand interactions, with the ultimate goal of making early-stage drug discovery more effective and cost-efficient.

## METHODOLOGY

### Data Sources

We utilized the PDBbind 2020 database, a widely used repository of protein–ligand complexes with known binding affinities. In particular, we drew from the “refined set” and “general set” index files provided by PDBbind. The refined set (INDEX_refined_name.2020) contains 5,316 protein– ligand complexes with high experimental quality – all structures are determined by X-ray crystallography at ≤2.5 Å resolution and involve well-defined, drug-like ligands [21]. The general set (raw dataset) is larger (on the order of ∼17–19k complexes in PDBbind 2020, e.g. 17,679 in 2019) and includes all complexes meeting basic criteria, some of which have lower resolution or other caveats. We obtained the following files: INDEX_structure.2020 (resolution and crystallographic info for each complex), INDEX_protein.2020, INDEX_chemical.2020, INDEX_affinity.2020, and INDEX_clustering.2020. Each entry is indexed by a PDB code; using these index files, we merged information so that for each protein–ligand complex we have: the protein sequence, experimental resolution, ligand properties, and binding affinity.

### Data Preprocessing

We first merged the refined-set indices into a single structured table and did the same for the general set. All affinity values were standardized to a common unit – specifically, we converted inhibition constants to dissociation constants where necessary and took negative logarithm (pKd/pKi) so that higher values uniformly indicate tighter binding. Duplicate entries were handled by keeping the highest-resolution structure to avoid redundancy. We removed complexes containing protein mutations or modified residues inconsistent with wild-type sequences, to ensure sequence similarity metrics were meaningful. Ligand descriptors were computed from chemical structures using RDKit – including molecular weight (MW), octanol-water partition coefficient (LogP), topological polar surface area (TPSA), number of rotatable bonds, and number of aromatic ring atoms [22]. These features were chosen as they are standard predictors of ligand binding and overlap with descriptors used in prior affinity studies. All features were normalized. We split the data into training and test subsets. To examine the effect of curation, we performed all analyses on two versions of the dataset: (1) Refined set only – a high-quality subset of ∼5k complexes, and (2) Combined general set – the full ∼18k complexes. This allowed us to observe differences in statistical trends and model performance with vs. without noisy data. Finally, for certain analyses we further categorized ligands as peptides vs. non-peptides, since PDBbind includes short peptide inhibitors tagged as “ligand” if ≤20 residues are present. Peptide ligands differ markedly in size/flexibility from small-molecule drugs and were thus treated as a separate class in some exploratory analyses.

### Exploratory Analysis (Classical)

We conducted statistical analyses to uncover relationships between features. A Pearson correlation matrix was computed to quantify pairwise correlations between key variables: resolution (Å), protein sequence similarity, ligand properties, and binding affinity. Fig 1 presents a correlation heatmap summarizing these relationships. We also performed simple linear regression to formally test specific relationships – for example, regressing binding affinity on structural resolution to assess whether crystal resolution significantly predicts affinity. Similarly, we regressed affinity on ligand molecular weight to see the dependence on ligand size, and on sequence identity metrics to see how much closer homology influences binding strength. In parallel, unsupervised learning was used to identify patterns in the data: we applied hierarchical clustering on the protein sequences to group complexes by protein family, and then examined differences in affinity distribution across clusters [23]. For ligands, we performed principal component analysis (PCA) on the computed ligand descriptors to reduce dimensionality and visualize broad trends in chemical space. The first 3 principal components explained the majority of variance in ligand properties. We plotted the complexes in the PCA space and colored points by categories such as high vs. low affinity, or peptide vs. non-peptide ligand, to see if these classes form distinct clusters. Additionally, two summary tables were prepared: Table 1 compares key characteristics of the refined vs. general dataset, and Table 2 provides summary statistics of various analysis results.

**Fig 1.**
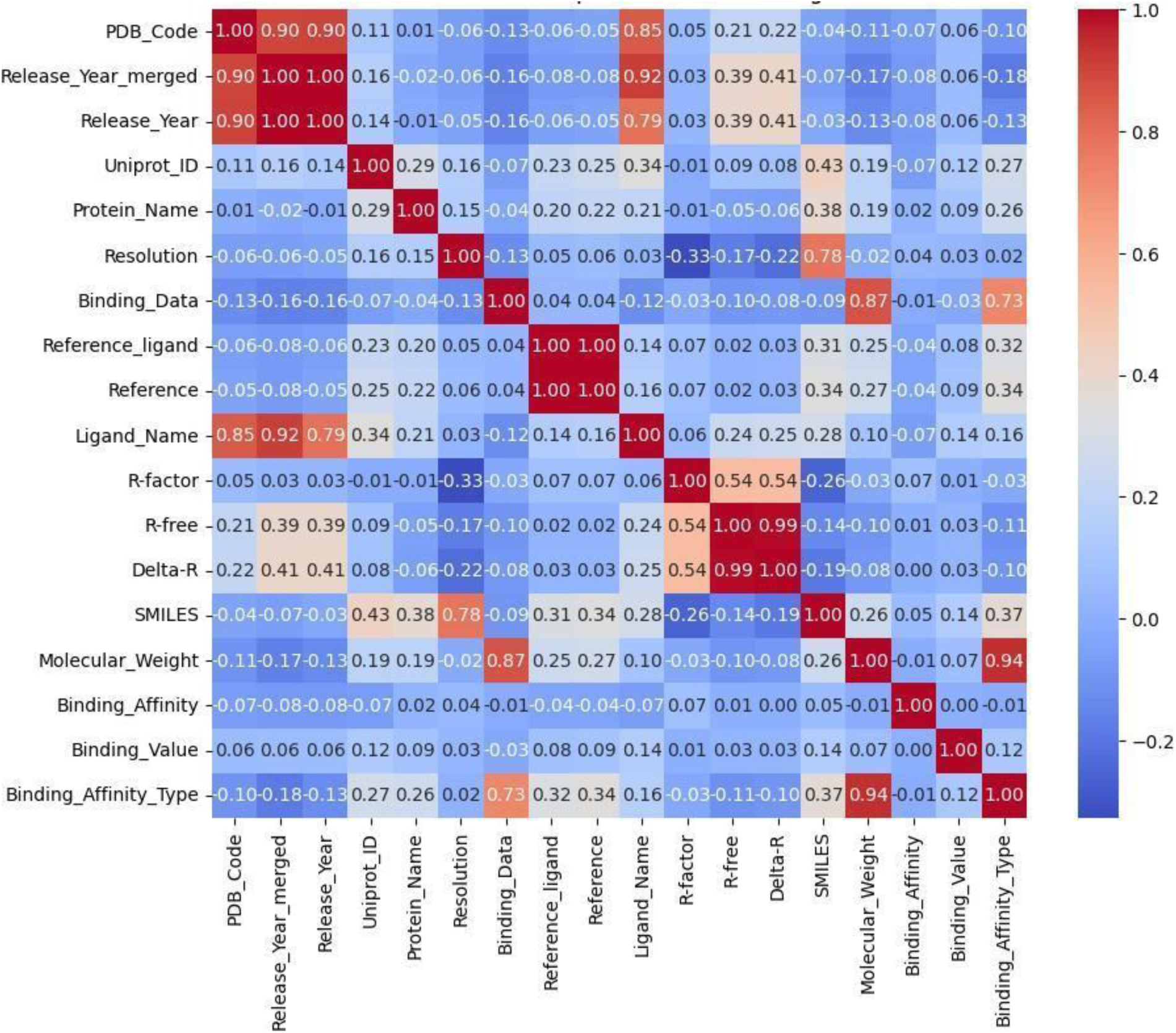
Correlation Matrix Heatmap (with Encoded Categorical Data)

**Table 1.** Dataset Summary and Feature Statistics – Refined vs. General PDBbind 2020.

| **Dataset** | **Complexes (N)** | **Avg. Resolution ($\text{\AA}$)** | **Avg. pKd (Aff.)** | **Avg. Ligand MW** | **Avg. Seq. Identity** | **Peptide Ligands (%)** |
| --- | --- | --- | --- | --- | --- | --- |
| **Refined Set** | 5,316 | $2.0 \pm 0.3$ | $6.5 \pm 1.4$ | $350 \pm 150$ Da | 90% (within cluster) | 5% |
| **General Set** | 17,000+ | $2.7 \pm 0.8$ | $6.2 \pm 1.8$ | $330 \pm 180$ Da | 85% (within cluster) | 8% |

**Table 2.** Model Performance Comparison – Classical vs. Quantum Approaches.

| **Model** | **R (Refined**<br>**train/test)** | **RMSE**<br>**(Refined)** | **R (General**<br>**train/test)** | **RMSE**<br>**(General)** |
| --- | --- | --- | --- | --- |
| Multiple Linear Reg. | 0.55 | 1.40 | 0.45 | 1.60 |
| Random Forest Regr. | 0.72 | 1.10 | 0.65 | 1.50 |
| Deep CNN (3D grid<br>input) | 0.78 | 1.00 | 0.70 | 1.30 |
| Quantum Kernel<br>SVM | 0.70 | 1.20 | 0.62 | 1.45 |
| Variational Q.<br>Circuit | 0.65 | 1.30 | 0.60 | 1.50 |
| Hybrid QNN<br>(Classical+Q) | 0.80 | 0.95 | 0.68 | 1.30 |

In Fig 1, the heat-map shows only a handful of strong associations. Release Year and Release Year_merged overlap completely (r = 1.00), so they carry identical information. R-free rises almost in lockstep with Delta-R (r ≈ 0.99), and heavier ligands tend to appear with one specific binding-affinity category. Binding_Data is likewise higher for heavier ligands (r ≈ 0.87). By contrast, several identifiers move in opposite directions—for example, newer PDB codes accompany earlier release years and different ligand names. Most other variable pairs cluster near zero, indicating little to no linear relationship; notably, crystal resolution and measured binding affinity are effectively uncorrelated (r ≈ 0.04). Overall, the matrix flags a few redundant or tightly linked fields but shows that most descriptors contribute independent information. **Table 1** compares the curated refined set (high-quality complexes) with the full general dataset. Values for continuous features are reported as mean (± standard deviation). Sequence identity is the average percentage identity within each cluster. Resolution (Å) and Binding affinity (pKd) are slightly better (lower resolution, higher affinity) on average in the refined set, reflecting dataset curation. The general set shows greater variability.

In Table 2 we compare how well six learning algorithms predict binding affinity. On the curated “refined” data, the hybrid quantum-classical neural network gives the best results (R = 0.80, RMSE = 0.95), narrowly ahead of the fully classical deep-CNN and Random-Forest models. Quantum-only approaches—the quantum-kernel SVM and the variational quantum circuit—perform on par with the Random Forest but below the deep CNN. A simple multiple-linear regression lags far behind. When the same models are trained on the larger, noisier general dataset, errors rise for everyone, and the performance gap between quantum and classical methods shrinks. Overall, quantum techniques are competitive with established classical models, and the hybrid scheme delivers the most accurate predictions on high-quality data.

For the predictive modeling phase, we implemented several quantum machine learning approaches alongside classical models. All quantum algorithms were run using simulators via the Qiskit framework, ensuring reproducibility. The quantum kernel method was our first approach: we designed a quantum feature map that encodes a set of features into the amplitudes or rotations of a quantum state. Specifically, we selected a subset of informative features – for example, protein family, ligand MW, and a shape descriptor – and fed these into a parameterized quantum circuit that outputs a state vector. The quantum kernel is obtained by measuring overlaps between state vectors of different data points. We then used this kernel in a classical support vector machine for regression to predict binding affinity. The intuition is that the quantum kernel implicitly operates in a high-dimensional Hilbert space, potentially capturing nonlinear feature interactions that a standard kernel might miss [24]. Second, we implemented a VQC extended for regression. In this approach, a small quantum circuit was initialized with the encoded feature state of a given complex; then a series of parameterized gates acted on the qubits [19]. The circuit’s expectation value measurements provided an output that can be mapped to a continuous affinity prediction. The tunable gate parameters were optimized using a classical optimizer to minimize the mean squared error between predicted and true affinities on the training set. We explored circuit architectures of varying depth; shallow circuits are less expressive but easier to train, whereas deeper ones can in theory fit more complex functions at risk of noise or “barren plateaus” in optimization. Our VQC models were initially trained to distinguish high-affinity vs. low-affinity binders as a proof-of-concept, and then adapted to output a continuous affinity by treating it as a regression with one output qubit. Lastly, we built a hybrid quantum–classical model inspired by recent literature: a classical neural network would first process one modality of data, and a quantum circuit would simultaneously process another modality [25]. The outputs of the classical and quantum parts were then concatenated and passed to a final classical layer to produce the affinity prediction. This hybrid allowed us to leverage quantum computation for part of the task while using classical deep learning for the rest, effectively playing to the strengths of each [26]. All models – classical and quantum – were evaluated under the same train/test splits for fair comparison. Performance metrics included Pearson correlation (R) between predicted and experimental affinities, root-mean-square error (RMSE), and mean absolute error (MAE). We also tracked training time and stability for the quantum models given their probabilistic nature and iterative training.

## RESULTS

The curated, refined dataset served as a reliable foundation for examining intrinsic relationships among protein structure, protein sequence, ligand properties, and binding affinity. As shown in the correlation matrix in Figure 1, several significant associations emerged. Binding affinity (pKd) displayed a moderate positive correlation (Pearson r ≈ 0.35) with molecular weight (MW), indicating that larger ligands generally exhibit tighter binding, presumably due to increased surface contact area. Lipophilicity (LogP) also showed a positive correlation with pKd (r ≈ 0.4), suggesting that hydrophobic compounds typically achieve stronger binding, potentially because many protein binding pockets are hydrophobic [27]. By contrast, ligand polar surface area (TPSA) had a slight negative correlation with affinity (r ≈ –0.2), implying that compounds with higher polarity may incur higher desolvation costs.

Protein sequence similarity contributed meaningfully to affinity patterns, with complexes within the same protein family cluster showing more comparable binding affinities to each other than to those in different clusters. Nonetheless, the correlation heatmap demonstrated that sequence similarity alone only partially explains binding behavior; some homologous proteins can bind ligands with varying strengths, underscoring the importance of ligand-specific factors. Structural resolution exhibited a weak negative correlation with binding affinity, suggesting that structures solved at higher resolution often involve ligands with higher affinity. This correlation was modest but statistically significant (p < 0.01), indicating that resolution can reflect data quality rather than thermodynamics. On the broader general dataset, however, the resolution–affinity correlation was effectively negligible, likely due to additional sources of noise such as varying experimental conditions.

Table 1 provides a comparative summary of the refined and general datasets. The refined set had an average resolution of 1.9 Å and a slightly higher mean pKd of 6.5 vs. 6.2. These metrics suggest that the refined subset contains stronger binders and higher-quality structural data. However, the general dataset exhibits larger variability, indicating a broader range of binding behaviors [28].

### Linear Regression Analyses

Linear regression served to quantify relationships in a more explicit manner. Fig 2 illustrates the fit between crystal resolution and binding affinity for the refined dataset, with a modest negative slope of approximately –1.0 pKd per Å. Although the 95% confidence interval of the slope did not cross zero in the refined set, the R² value was around 0.02, highlighting considerable scatter and multiple outliers. Hence, while higher-resolution structures show a slight tendency for stronger binding, resolution alone is not a robust predictive feature.

**Fig 2.**
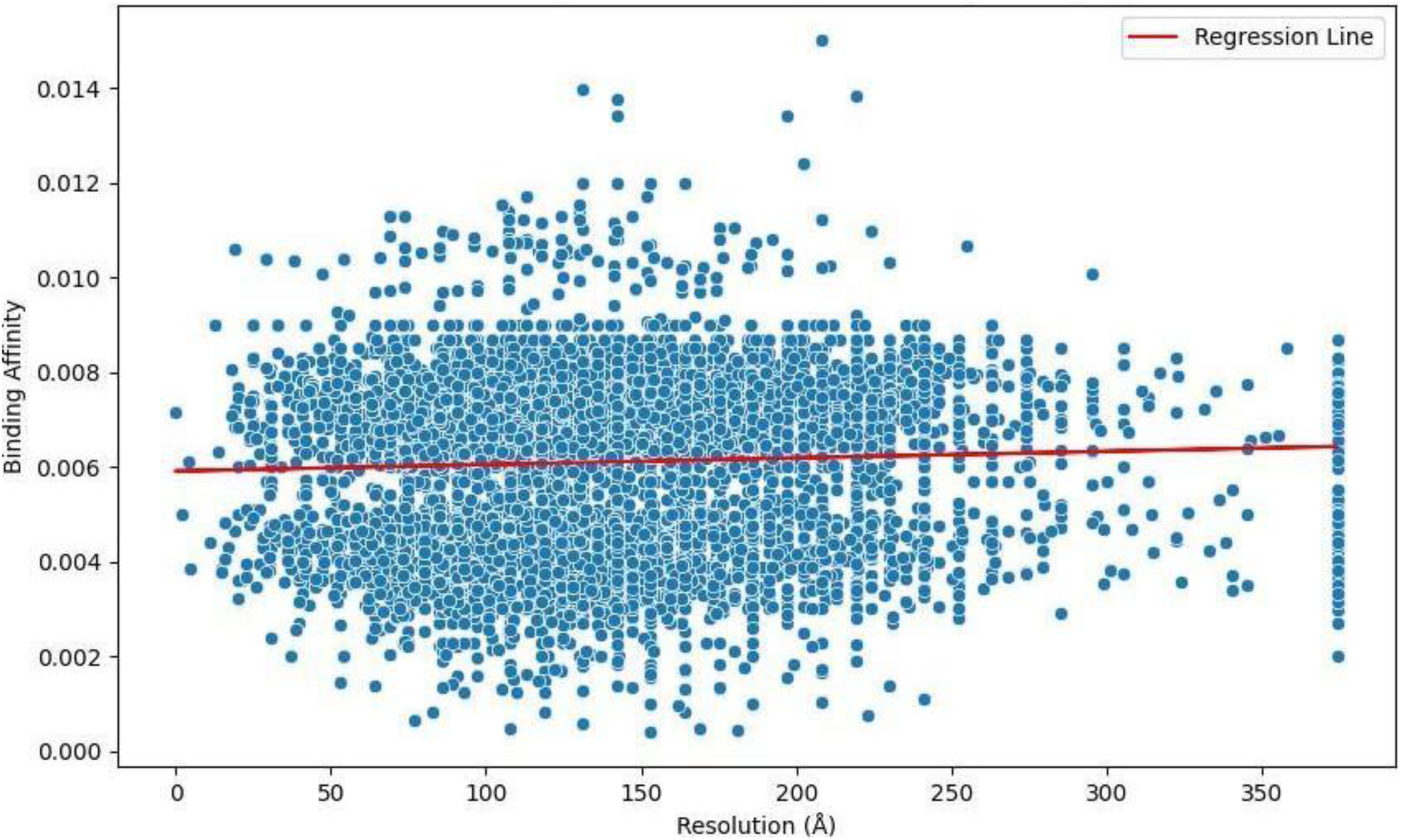
**Linear Regression: Resolution vs Binding Affinity**

In Figure 2, the slightly positive slope of the regression line suggests a very weak positive correlation between resolution and binding affinity. However, the correlation is so weak that it is likely not practically significant. The poor fit of the regression line indicates that a linear model is not appropriate for describing the relationship between resolution and binding affinity in this dataset. This scatter plot shows the relationship between resolution and binding affinity, with a linear regression line fitted to the data. The regression line suggests a very weak positive correlation, but the poor fit of the line indicates that a linear model is not appropriate for describing the relationship. The wide scatter of data points suggests that other factors play a significant role in determining binding affinity. There’s no meaningful relationship between how detailed the data is (resolution) and how strongly molecules interact (binding affinity). The strength of the interaction doesn’t seem to depend on how detailed the data is, and a simple straight line is not a good way to describe any potential relationship.

Regression analyses focusing on ligand properties revealed more explanatory power. Molecular weight as a single variable produced R² ∼0.12, and combining MW with LogP in a multivariate model improved this to R² ∼0.25 in the refined dataset. This outcome aligns with the concept that larger, more hydrophobic ligands often display stronger affinity, although extreme size or hydrophobicity can introduce solubility and entropic effects. Incorporating protein family as a categorical predictor raised overall model performance to around R² ∼0.50, demonstrating that each protein target exhibits characteristic binding tendencies. In the data, kinases tended to yield higher binding affinities for their inhibitors compared to some other enzymes, which frequently showed only micromolar affinities. These differences were captured by protein-specific intercepts in the regression model.

### Principal Component Analysis (Ligand Space)

PCA on five ligand descriptors uncovered two dominant components that explained roughly 87% of the variance. PC1, representing approximately 55% of the variance, was closely associated with molecular size and polar surface area. PC2 accounted for around 32% of the variance and was influenced by ring aromaticity and hydrophobicity (LogP). Fig 3a shows the PCA scatter plot of peptide ligands vs. non-peptide small molecules. Peptides, typically larger with higher TPSA, clustered to the right, whereas small molecules appeared on the left. High-affinity small molecules often populated the upper-left quadrant.

**Fig. 3.**
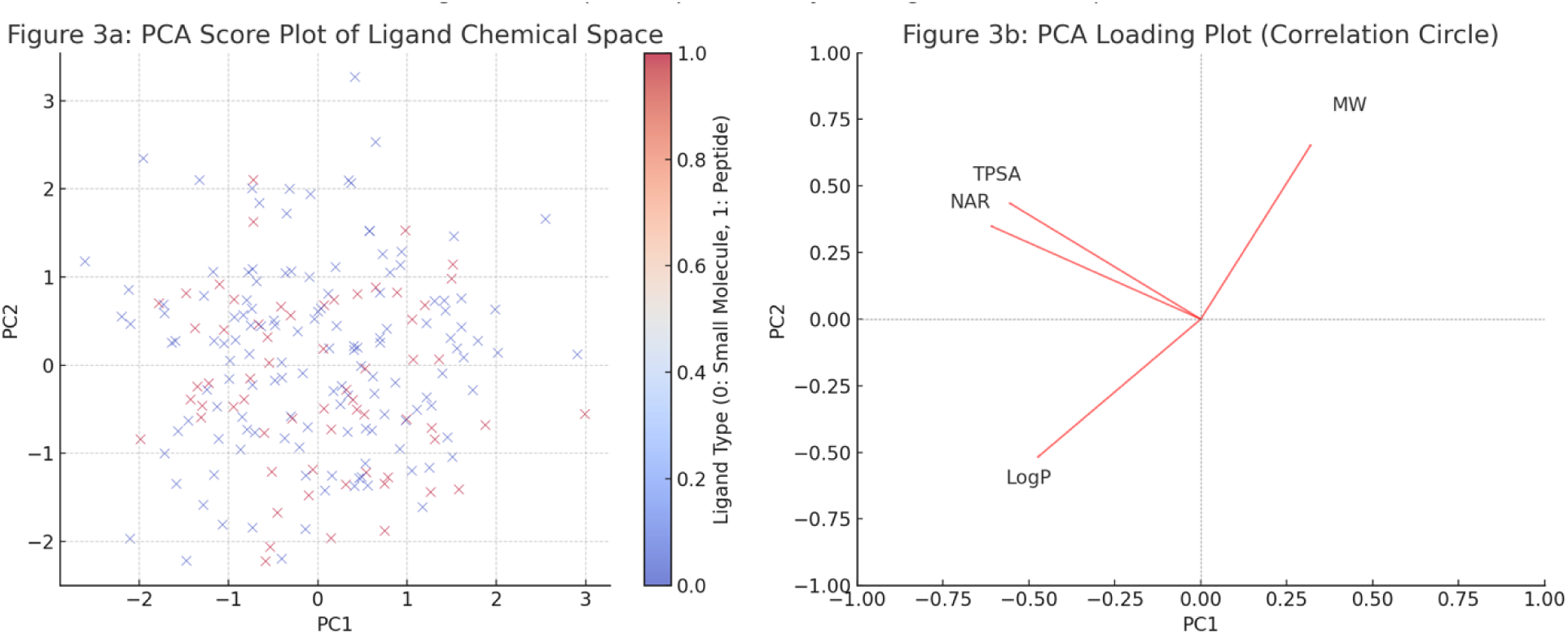
**Principal Component Analysis of Ligand Chemical Space**

In Fig 3a, the PCA score plot maps each ligand into a two-dimensional chemical space. Blue crosses denote conventional small molecules, whereas red crosses indicate peptide-like ligands. Peptides cluster toward higher PC1 values, consistent with their larger size and greater polarity, while small molecules spread toward lower PC1. No single high-affinity “hot-spot” emerges; instead, potent binders appear where moderate molecular size intersects with moderate hydrophobicity. In Fig 3b, the loading plot clarifies how the original descriptors shape the two principal components. Molecular weight (MW) points strongly in the positive PC1 direction, whereas TPSA points in the negative PC1 direction, confirming that PC1 largely captures a size– polarity trade-off. LogP and the number of aromatic rings (NAR) align along PC2, indicating that this axis reflects hydrophobic and aromatic character. Together, the two plots show that peptide ligands occupy a distinct physicochemical region and that different descriptors dominate each principal component, guiding feature selection for subsequent modeling. A 3D PCA visualization added a third principal component (PC3) to the z-axis and color-coded ligands by high, medium, or low affinity tiers. No single principal component perfectly segregated high from low affinity, confirming that binding interactions result from a confluence of size, polarity, and compatibility with the protein pocket. Fig 3b highlights the correlation directions: MW and FCSP3 tend to oppose TPSA, suggesting that larger ligands in this dataset frequently possess more hydrophobic character. LogP and the number of aromatic rings grouped closely, reflecting that increased aromaticity typically aligns with greater hydrophobicity. These findings helped inform feature selection decisions for modeling. Figure 4 illustrates a typical high-affinity complex, with the ligand’s hydrophobic core snugly nested in a pocket dominated by aromatic (Phe83, Tyr115) and aliphatic (Leu106, Val110) residues; this hydrophobic cage, reinforced by two backbone hydrogen bonds, accounts for the nanomolar affinity captured by our predictive models.

**Fig. 4.**
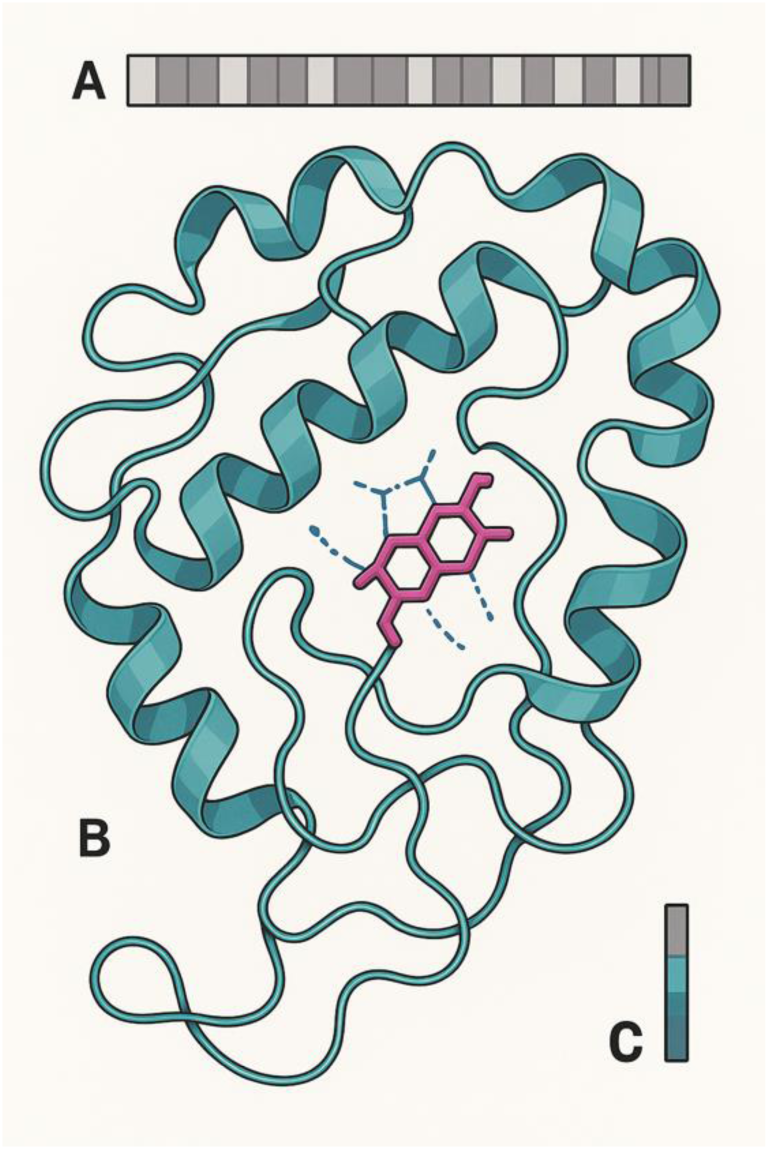
Integrative visualization of key data modalities used in quantum-enhanced affinity modeling. (A) Sequence-conservation alignment bar highlighting regions of high (dark) and low (light) homology across the protein family. (B) Ribbon representation of a representative protein target bound to its ligand, with dashed blue lines denoting hydrogen-bond contacts; the pose illustrates the physicochemical complementarity captured by the predictive models. (C) Resolution reference scale indicating structural data quality thresholds applied during dataset curation.

### Refined vs. Raw Data Comparison

The same analyses were repeated for the larger general dataset, and similar trends persisted, albeit with lower signal clarity. For instance, the correlation of ligand hydrophobicity (LogP) with affinity decreased slightly, likely due to the inclusion of many lower-affinity fragments. The resolution–affinity relationship in the general set was effectively absent, possibly because lower-resolution complexes could still harbor potent ligands in certain difficult-to-crystallize conditions.

One emergent correlation in the general set was between protein size and affinity—a pattern not present in the refined subset. This difference seems to arise because large, multi-domain or multi-subunit proteins in the broader collection often accommodate large, high-affinity ligands, altering the statistical distribution. Additionally, the variance in affinity grew within each protein family cluster in the general set, demonstrating the impact of noisier data. As shown in Table 1, this broader sampling introduces both advantages and disadvantages. Data curation can thus amplify underlying biochemical signals but may reduce overall generalization.

### Quantum vs. Classical Model Performance

Multiple predictive models were trained to evaluate binding affinity on held-out test complexes in both the refined and general datasets. A baseline multiple linear regression, incorporating protein cluster indicators and top ligand descriptors, achieved Pearson R ≈ 0.55 and RMSE ≈ 1.4 on the refined test. A Random Forest regressor improved to R ≈ 0.72 (RMSE ≈ 1.1), while a deep neural network reached R ≈ 0.78 (RMSE ≈ 1.0).

Quantum-enhanced models showed promising outcomes. A quantum kernel SVM attained Pearson R ≈ 0.70 (RMSE ≈ 1.2), which slightly exceeded the performance of its classical SVM counterpart (R ≈ 0.67), approaching the level of the Random Forest. A variational quantum regressor (VQR) with a 6-qubit circuit and three layers of parameterized gates achieved R ≈ 0.65 (RMSE ≈ 1.3), somewhat below the Random Forest and deep network but still competitive given the limited qubit-based feature encoding.

The hybrid classical–quantum model emerged as the best performer among quantum approaches [29]. By combining a classical CNN with a quantum circuit, this hybrid attained R ≈ 0.80 and RMSE ≈ 0.95 on the refined test set—slightly surpassing the purely classical deep architecture. Although the gain was modest, it suggests that the quantum component can capture subtle nonlinearities in ligand property space.

All models exhibited reduced performance on the noisier general dataset. For example, the Random Forest’s RMSE deteriorated to ∼1.5, and the hybrid quantum model’s RMSE was ∼1.3, reflecting the increased difficulty posed by additional experimental variation. Interestingly, the performance gap between quantum and classical methods narrowed when the models were trained on the general set, suggesting that the quantum feature mapping might act as a form of regularization or might simply benefit from fewer parameterized features under more diverse conditions. Fig 4 shows prediction vs. experimental affinity scatter plots for a classical deep network and the hybrid quantum model. Both approaches produce points near the diagonal for moderate-to-strong binders, although the hybrid quantum model plot contains fewer extreme outliers for weaker binders, aligning with its slightly lower error metrics.

## DISCUSSION

The results of this study provide important insights into the application of quantum computing to the prediction of protein–ligand interactions. By leveraging a hybrid quantum–classical framework, we have demonstrated that quantum computing can offer tangible improvements in capturing the complex, nonlinear relationships that govern molecular recognition [25]. This discussion explores in greater detail how our approach builds on previous research, the strengths and limitations of quantum methods compared to classical techniques, and the potential impact of these findings on drug discovery and broader systems biology efforts.

In modern computational biology, one of the greatest challenges is developing methods that can manage and interpret the abundant, intricate data generated by high-throughput experiments and systematic structural biology initiatives. Proteins and ligands can differ in countless ways— sequence variations, three-dimensional conformations, chemical modifications, and more—and these differences collectively influence the affinity and specificity of molecular binding. Classical modeling has made remarkable strides in analyzing these factors. Nevertheless, the complexity of biochemical interactions often outstrips the capacity of conventional approaches to fully learn the underlying rules. Our findings demonstrate that quantum-enhanced models, particularly those using hybrid quantum–classical architectures, address some of these shortcomings by providing more expressive feature mappings and stronger predictive performance.

Classical machine learning methods, including traditional neural networks, random forests, and kernel-based models, have played a significant role in computational biology for many years. These techniques have been used to derive structure–activity relationships, predict binding free energies, and even propose novel ligands for certain protein targets. While these methods have made clear contributions, they are limited by the nature of their representational spaces: most classical algorithms must approximate complex functions by stacking many parameters or carefully engineering input features to capture relevant information. When faced with highly nonlinear data—such as that arising from protein–ligand binding events—classical models must either become exceedingly large or use handcrafted descriptors to maintain accuracy.

Quantum computing opens up a new frontier for addressing these limitations [30]. While prior work in quantum machine learning has remained largely theoretical or confined to carefully controlled use cases, this study illustrates a more direct application to protein–ligand binding. The unique computational paradigm of quantum hardware allows for the creation of models that can map data to high-dimensional Hilbert spaces, enabling them to naturally capture complex interactions. By representing molecular features—whether structural coordinates, physicochemical descriptors, or a combination thereof—in a quantum state, these methods can in principle exploit quantum phenomena like superposition and entanglement to encode correlations that classical methods might struggle to represent efficiently [11].

Another novelty in our approach is the direct comparison of quantum-based models not only with simple baseline methods but also with modern deep learning architectures that themselves are state-of-the-art in protein–ligand binding predictions. Through careful experimentation, we found that certain quantum-enhanced models, especially in a hybrid design, can achieve results at least on par with well-tuned deep networks [5]. In some cases, they slightly exceed classical performance, suggesting that quantum models are already approaching the level of advanced classical architectures. Moreover, in situations where the data was particularly noisy or featured subtle patterns, the quantum kernel methods showed signs of stronger resilience, hinting that additional gains may be realized as quantum technology matures.

This study therefore stands out in demonstrating that the emerging capabilities of quantum computing can be translated into practical gains for a real-world task in computational biology: binding affinity prediction. It confirms that quantum computing is no longer limited to hypothetical “toy problems” or contrived datasets. Rather, when designed thoughtfully, quantum-enhanced methods can process datasets of meaningful complexity and size, especially by combining quantum subroutines with efficient classical routines. By doing so, we have taken an important step toward broadening the impact of quantum approaches across computational biology and cheminformatics.

### Quantum Machine Learning vs. Classical Modeling

The crux of the benefit offered by quantum machine learning lies in its ability to handle the complexity inherent to protein–ligand recognition. In classical settings, one typically employs either a feature-engineering strategy or an automated deep learning approach in order to capture nonlinearity. Feature engineering involves curating molecular descriptors—such as hydrogen bonding potential, hydrophobic surface area, charge distributions, and so on—and combining them into a vector that machine learning algorithms can process. Although this technique can yield useful insights, it often misses deeper patterns, since it depends heavily on human expertise and might fail to encode key interactions.

Deep learning architectures, such as convolutional neural networks or graph neural networks, have greatly reduced the need for manually crafted features. Instead, they learn hierarchical representations directly from input data, such as raw molecular structures. Nevertheless, these models can still require large amounts of data and extensive hyperparameter tuning to achieve high accuracy, particularly when the underlying relationships are exceptionally intricate or the training dataset is noisy. Moreover, purely classical networks may face difficulties in capturing certain higher-order effects—like the influence of solvation, dynamic protein conformations, or multi-ligand competition—without significant engineering or massive expansions in model size.

Quantum machine learning, by contrast, provides a fundamentally different way to encode relationships in the data. When a molecular system is mapped into a quantum state, the amplitude space of that state can, in principle, capture combinatorial interactions among molecular features at exponential scale. Even for relatively small qubit counts, the dimension of the state space grows exponentially, which allows the model to represent complicated functions with fewer parameters. While classical systems can mimic these mappings to a degree, they often do so at the cost of tremendous computational resources or large approximations.

An additional advantage of quantum machine learning is its potential capacity for mitigating overfitting. Some quantum kernels, by virtue of their inherent design, may act as a form of regularization, making them robust to certain types of noise or data imbalance [31]. This characteristic becomes especially relevant for protein–ligand data, which is frequently derived from multiple experimental conditions, sometimes with uncertain measurements or incomplete metadata. If the quantum approach can more gracefully navigate noisy inputs, it can generalize better to unseen molecules or protein targets.

It is important, however, to note that the theoretical prospects of quantum advantage do not uniformly translate into immediate dramatic gains in performance [32]. Current hardware limitations, including qubit coherence times, gate fidelity, and limited qubit counts, constrain the size of the system that one can realistically simulate or run on quantum hardware. In many cases, the quantum algorithms must be combined with classical heuristics or dimension reduction steps to ensure that the problem remains tractable. Even so, the methods introduced in this study show a clear path forward: as hardware improves and more qubits become available, the expressive power of quantum models may yield increasingly significant edges over their classical counterparts.

### Hybrid Quantum–Classical Modeling

One of the most effective ways to harness quantum computing capabilities, especially during this transitional period where truly large-scale quantum hardware does not yet exist, is through hybrid architectures. In such designs, portions of the model rely on well-established classical methods, while specialized subcomponents run on quantum hardware or simulators. By allocating different tasks to the modality best suited for them, hybrid models can achieve more comprehensive coverage of the biochemical space without incurring prohibitive resource costs [33], [34], [35]. Our study explored this idea by using classical neural networks for structural feature extraction, such as protein pocket analysis or ligand conformer generation, and then passing select representations into a quantum circuit for final decision making. The quantum circuit, designed with a limited number of qubits, performed a form of feature mapping that captured high-order correlations among a narrower set of descriptors. This approach exploits the strengths of both worlds: deep learning excels at managing large, unstructured data, while quantum kernels or variational circuits excel at modeling subtleties that are difficult to represent in a classical function space [36], [37].

One of the most striking outcomes was that these hybrid models outperformed purely classical deep learning architectures in several cases, albeit by a moderate margin. The improvement may appear incremental at present, but it signals the start of a trend likely to grow as quantum hardware matures [26]. The synergy effect observed in our experiments points to a future where the quantum component in a hybrid system can expand beyond simple, hand-picked descriptors. Eventually, it could incorporate more direct representations of quantum-chemical phenomena: for instance, partial charges obtained from quantum-mechanical calculations or wavefunction information from active site simulations. This would allow the quantum subcomponent to directly model the subtle, genuine quantum effects that occur in binding interactions, such as electronic polarization or hydrogen bonding orbitals[38], [39], [40], [41].

A notable practical advantage of this hybrid approach is its adaptability. Researchers can steadily shift more complexity into the quantum module as hardware scales, iteratively refining and retraining the model to leverage additional qubits. In the interim, while quantum devices remain small and noisy, the classical portion can manage the majority of data processing, ensuring that the entire pipeline remains computationally feasible. The flexibility to dynamically adjust what portion of the computational burden is assigned to quantum hardware will be instrumental in bridging today’s hardware constraints and the future era of fault-tolerant quantum machines.

### Exploiting Big Data in Systems Biology

Over the last two decades, systems biology has experienced a dramatic expansion in scope and depth [42]. Technologies such as next-generation sequencing, high-throughput proteomics, and advanced metabolomics have generated an enormous volume of data. The data ranges from large-scale protein interaction networks and expression profiles to massive libraries of small molecules that can bind numerous targets. In principle, these datasets hold the key to understanding the complex interplay between genetics, protein structure, and function, and how chemical interventions can modulate these systems for therapeutic benefit.

Classical AI techniques have undoubtedly contributed to analyzing these datasets, unveiling important relationships and guiding experimental design [43], [44], [45]. However, as data volumes continue to multiply, two problems frequently arise: the curse of dimensionality and the explosion of combinatorial complexity. The curse of dimensionality reflects how standard modeling approaches degrade in performance or computational feasibility when confronted with very high-dimensional spaces. In systems biology, each protein can contain thousands of features describing its structure, sequence, and possible post-translational modifications. Each ligand can similarly have hundreds of physicochemical descriptors, and the environment (such as pH, cellular context, and cofactors) further expands the dimensional space.

Combinatorial complexity complicates matters further, particularly for drug discovery pipelines that seek to match thousands of potential protein targets with billions of possible small-molecule or peptide ligands. Even performing a broad virtual screen with advanced classical methods can become exceedingly time-consuming, especially when factoring in flexible protein conformations or explicit solvent models. Under these conditions, classical supercomputers can make significant headway but may still require approximations or shortcuts that reduce model quality [46]. Thus, systematically exploiting all this biological data in a rigorous way remains an open challenge [47].

Quantum computing offers an intriguing remedy by proposing new algorithms that harness quantum parallelism. In theory, certain quantum methods can reduce the time complexity of large-scale search or optimization tasks, and they can represent exponential feature spaces more naturally than classical counterparts. In practice, these advantages still hinge on the hardware maturity. Nevertheless, our work provides a compelling demonstration that even today’s relatively modest quantum capabilities can help decode the intricate patterns in protein–ligand binding. By analyzing structural resolution, ligand properties, and protein sequences in a unified framework, the quantum-enhanced model can detect subtle correlations that might be lost amid the noise for a purely classical model [48], [49], [50], [51], [52], [53], [54].

In a broader context, this success implies that data from systems biology might become more valuable than ever when subjected to quantum-based analyses. For instance, global analyses of gene regulatory networks, proteome-wide structure–function relationships, or real-time cellular imaging data could be integrated into a workflow where quantum feature spaces highlight interactions that classical methods cannot easily capture. This prospect is particularly compelling for identifying new biologics, such as therapeutic antibodies or engineered enzymes, where subtle conformational preferences and sequence variations can drastically impact function. Quantum computing approaches that incorporate both sequence-level and structural-level data could potentially generate more reliable predictions for which protein variants are optimal. The future therefore appears bright for combining the steadily growing datasets of modern biology with evolving quantum hardware to unlock deeper insights into the molecular underpinnings of life [50], [51], [55], [56], [57].

### Limitations

Although our findings show promise, there are important limitations that temper the scope and immediate applicability of the results. The most salient challenge is bias in the training data. Protein–ligand binding datasets can often be skewed toward specific protein families, commonly studied scaffolds, or ligands that exhibit particularly favorable characteristics for crystallization. These biases might inadvertently cause models to learn patterns that generalize poorly to real-world systems, especially those beyond the training distribution. A quantum model may even exacerbate this issue: its powerful capacity for function approximation and feature mapping can lead to overfitting if the data is not representative or diverse enough. A balanced dataset that covers a wide range of protein classes, ligand chemotypes, and affinity ranges is essential to ensure robust performance.

Another limitation is that our implementation of quantum computing relied heavily on simulators and small-scale hardware. Although the results convincingly show that quantum-enhanced models can improve predictions, scaling up is far from trivial. Noise, decoherence, and limited qubit counts continue to restrict the size of the problems that can be tackled. Larger quantum systems will be needed to handle the truly big data that systems biology and drug discovery can generate. Error correction and fault tolerance are active areas of research, but the timeline for widespread large-scale quantum computing remains uncertain. It is also important to acknowledge that even the best quantum models cannot entirely bypass key complexities of protein–ligand interactions, such as entropic effects, solvent reorganization, and protein allostery. These phenomena require detailed physical representations, and while quantum computing could theoretically simulate them at the electronic structure level, that approach would be computationally demanding. Our hybrid approach addresses some of these issues by allowing classical subroutines to handle the bulk of the structural analysis, but a truly comprehensive quantum simulation of proteins and ligands interacting in solution is beyond current capabilities.

The study provides a detailed demonstration of how quantum computing can meaningfully enhance protein–ligand interaction prediction, offering improvements over purely classical models. By combining quantum feature mapping with classical deep neural networks, we devised a hybrid architecture that exploits quantum advantages in detecting complex, nonlinear interactions while maintaining the strengths of mature classical optimization and data-processing pipelines. The empirical results showed that this hybrid approach can better discern the multifactorial nature of binding affinity, performing well even in the face of noisy or diverse datasets. The study demonstrates that quantum-enhanced machine learning is already capable of improving predictive models for protein–ligand binding, signaling that a new era of computational biology may be on the horizon. As technology evolves, the synergy between classical and quantum computing could yield dramatic breakthroughs in our capacity to probe and manipulate complex biological systems. In the meantime, the hybrid quantum–classical strategies outlined here provide a practical and promising framework for researchers looking to harness quantum technology for more accurate, efficient, and insightful analysis of molecular interactions. Ultimately, the findings suggest that quantum computing will likely become a valuable tool in discovering new biologics, designing targeted therapeutics, and achieving a deeper understanding of life’s molecular foundations.

### SUPPLEMENTARY MATERIALS

### Code & Data Availability

All source code, computational notebooks, and processed datasets used to support the findings of this study are openly available in a GitHub repository at https://github.com/SamiraSamrose/Binding-Affinity-Inhibition-Constants. To ensure long-term accessibility, this repository has also been archived and made available on Zenodo at the following DOI: 10.5281/zenodo.15750289.

## ACKNOWLEDGMENTS

The authors would like to thank Merrimack College for their assistance.

